# Cognitive Fitness in Ageing (COFITAGE): A Multimodal and Longitudinal Neuroimaging Dataset

**DOI:** 10.64898/2026.08.24.746635

**Authors:** Antoine Jacquemin, Jiqing Huang, Nikita Beliy, Christian Degueldre, Fraçois Meyer, Daphne Chylinski, Justinas Narbutas, Maxime Van Egroo, Eric Salmon, Puneet Talwar, Fabienne Collette, Gilles Vandewalle, Christine Bastin, Mohamed Ali Bahri, Christophe Phillips

## Abstract

**Purpose:** Brain ageing involves interrelated changes in molecular processes, neuroinflammatory mechanisms, brain macro- and microstructure, sleep physiology, and cognition. The 50 to 70 years age range represents a critical transition period, in which these subtle alterations may precede measurable cognitive decline and the onset of clinical neurodegenerative disease. To allow systematic investigation of these early alterations and the subsequent progression in brain aging, we provide an open-access data resource from a multidisciplinary longitudinal study integrating neuroimaging, genetics, sleep, and neuropsychological phenotyping with assessments at baseline and at 2-year follow-up.

**Acquisition and Validation Methods:** The baseline cohort comprises 101 community-dwelling participants (50-69 years old) who underwent magnetic resonance imaging (MRI) using a 3T protocol that included high-resolution structural imaging (T1- and T2-weighted), quantitative multi-parametric acquisitions with B1 mapping, and multi-shell diffusion-weighted imaging. Moreover, positron emission tomography (PET) imaging was performed using [18F]Flutemetamol or [18F]Florbetapir (amyloid-*β* tracers) in all participants, with a subset also undergoing [18F]THK-5351 PET (tau-related/neuroinflammation). The dataset was complemented by extensive phenotypic data, including sleep and neuropsychological assessments, and by genotype data through genome-wide analysis. 66 participants underwent a 2-year cognitive follow-up, enabling longitudinal analyses of cognitive trajectories. Data acquisition and curation were performed using standardized procedures, with systematic quality control to support reliable cross-sectional and longitudinal analyses.

**Data Format and Usage Notes:** All data are distributed in a BIDS-compliant format, and released in open-access (EBRAINS).

**Potential Applications:** This dataset supports multimodal analyses, allowing the identification of interpretable patterns characterizing brain ageing from multiple perspectives. It enables the comparison of different models to derive (semi)quantitative MRI parameters, the discovery of imaging biomarkers associated with early cognitive decline, and the monitoring or prediction of brain ageing progression. In addition, it offers focused coverage of adults aged 50–70 years, which is often underrepresented in existing healthy subjects public datasets.

**Key Points:**

- COFITAGE is a deeply phenotyped, longitudinal, multimodal dataset of 101 healthy late middle-aged adults (50-70 years).
- The COFITAGE dataset combines PET, MRI, sleep phenotyping, genotyping, and extensive neuropsychological assessment.
- The dataset supports diverse applications, from preclinical AD biomarker studies to investigations of hippocampal vulnerability, amyloid-tau-metabolism interactions, sleep-dependent modulation of molecular pathology, and multimodal predictive modeling of cognitive trajectories.

## I. Introduction

Understanding the early mechanisms of brain ageing remains a major challenge, as pathological processes associated with neurodegenerative diseases such as Alzheimer’s disease (AD) begin decades before the onset of clinical symptoms^1,2^. In midlife and early late adulthood, subtle alterations can already be detected across multiple biological levels, even in cognitively healthy individuals. At the molecular level, positron emission tomography (PET) studies have demonstrated that amyloid-*β* and tau accumulations begin years before measurable cognitive decline and are markers of preclinical AD^2,3^. At the structural level, magnetic resonance imaging (MRI) reveals early changes such as hippocampal atrophy and cortical thinning, which are associated with reduced memory performance, progressive memory decline, and an increased risk of subsequent cognitive impairment and neurodegenerative disease^1,4^. Beyond macrostructure, advanced MRI techniques provide access to microstructural properties of brain tissue, offering insights into processes such as myelin degeneration and axonal integrity that may precede volumetric changes ^5^.

Physiological and genetic factors further contribute to these early alterations. Sleep plays a fundamental role in memory consolidation and metabolic clearance, and disruptions in sleep architecture have been linked to increased amyloid and tau protein burden and cognitive decline^6,7,8^. In parallel, genetic factors, such as the presence of the apolipoprotein E (*APOE*) *ε*4 allele, modulate individual vulnerability to AD pathology and influence both brain structure and molecular accumulation^1,9^. These biological alterations ultimately manifest at the cognitive level, affecting domains such as episodic memory, attention, and executive functions, which are vulnerable to early decline due to AD^1,4^.

Importantly, these processes do not occur in isolation but interact dynamically across scales, contributing to inter-individual variability in ageing trajectories^1,4^. As a result, a comprehensive understanding of brain ageing requires the integration of molecular, structural, physiological, genetic, and cognitive measures within the same individuals.

However, few open-access resources combine positron emission tomography (PET), magnetic resonance imaging (MRI), sleep phenotyping, genotyping, and extensive neuropsychological assessment within the same individuals. The “Cognitive Fitness in Aging” (COFITAGE) project is a prospective, multimodal, longitudinal research initiative designed to characterize brain ageing in cognitively healthy late middle-aged adults. It was developed to address the lack of integrative datasets by combining molecular imaging of amyloid and tau, advanced structural and microstructural MRI, detailed assessments of sleep physiology, genetic information and extensive longitudinal cognitive phenotyping. This rich and multidimensional framework enables the investigation of interactions between these factors and their contribution to individual differences in cognitive trajectories. Several studies have already leveraged the COFITAGE dataset to examine the relationships between sleep^10,11,12^, molecular biomarkers^13,14,15,16,17^, hippocampal volume^18^ and cognition^19,20,21,22,23^.

The primary objective of this dataset paper is to present a comprehensive multimodal and phenotyped dataset acquired in cognitively healthy late middle-aged adults, featuring longitudinal data obtained through a two-year follow-up. The second objective is to provide harmonized PET, MRI, and phenotypic data organized according to the Brain Imaging Data Structure (BIDS) standard, thereby promoting reproducibility, transparency, and interoperability across studies^24,25^. The third objective is to enable the progressive release of advanced derivatives and to expand the dataset with additional modalities, including electroencephalography, transcranial magnetic stimulation, genetics and polysomnography, and extended longitudinal follow-up assessments at seven years.

## II. Methods

### II.A. Experimental Design

The study design is shown in Table 1, and the timeline of data acquisitions is provided in Figure 1. Participants first underwent a two-step screening procedure to ensure eligibility. The initial screening session included a brief neuropsychological evaluation and comprehensive questionnaires assessing medical history, including prior brain trauma, neurological and psychiatric disorders, as well as lifestyle factors such as caffeine and alcohol consumption, medication and drug use. Body mass index (BMI) was also objectively measured. This was followed by an in-laboratory overnight sleep assessment under full polysomnography (PSG) to identify potential sleep disorders, with a focus on sleep apnea and periodic limb movements (PLMs).

**Table 1:**
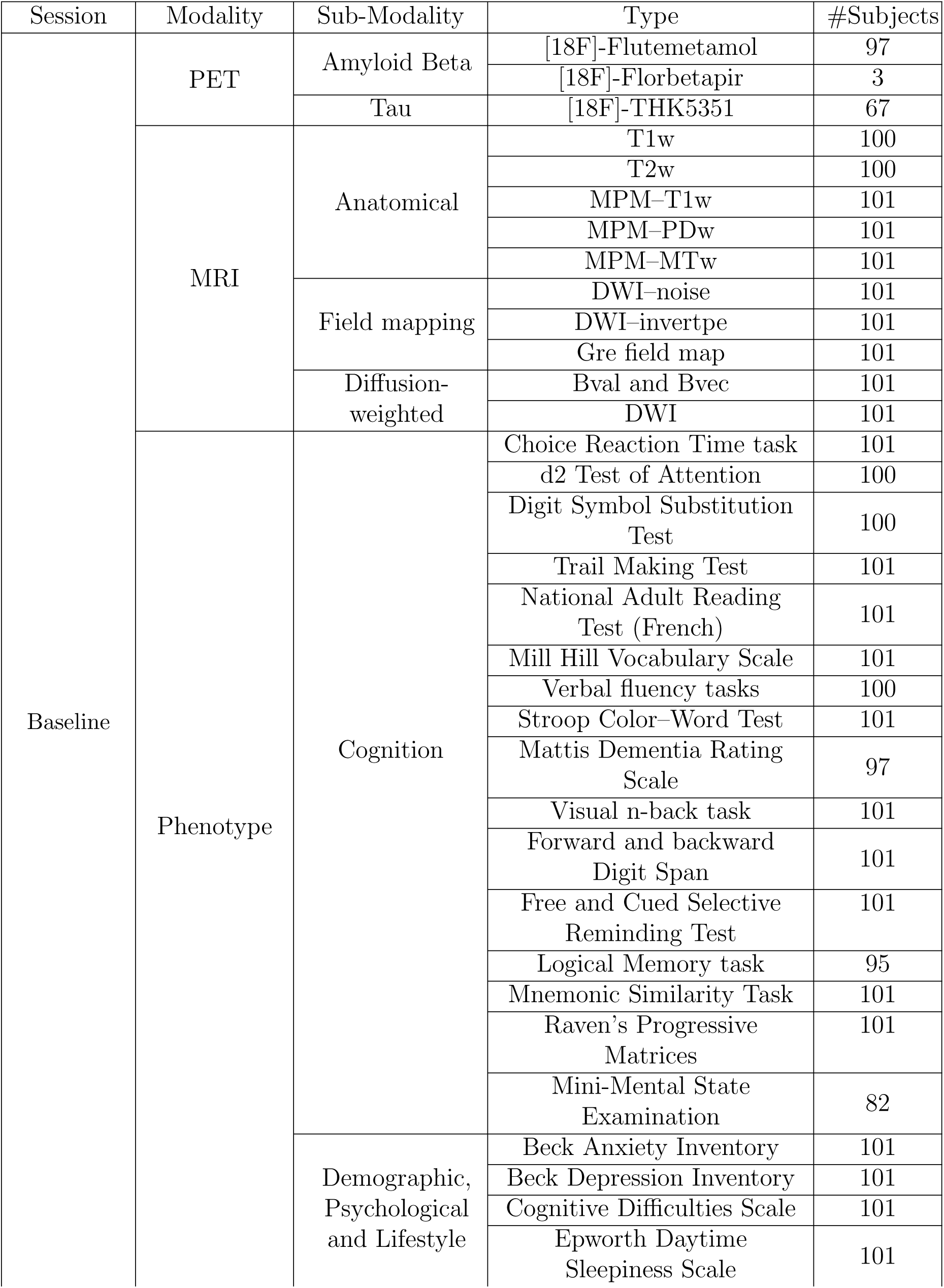
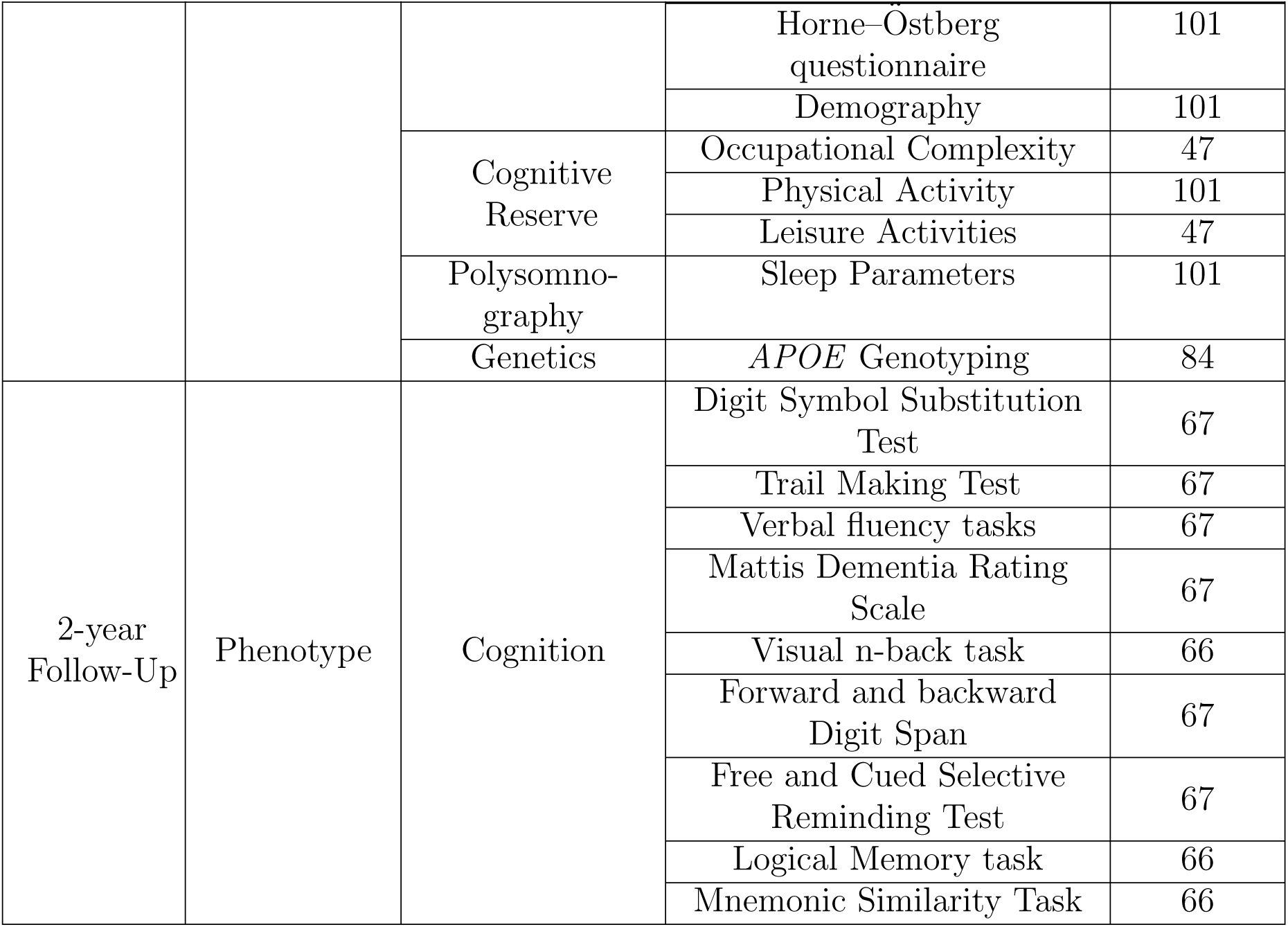
Schematic overview of the study design, illustrating the data available across sessions and modalities. The reported sample size reflects the data available among the 101 participants assessed at baseline.

**Figure 1:**
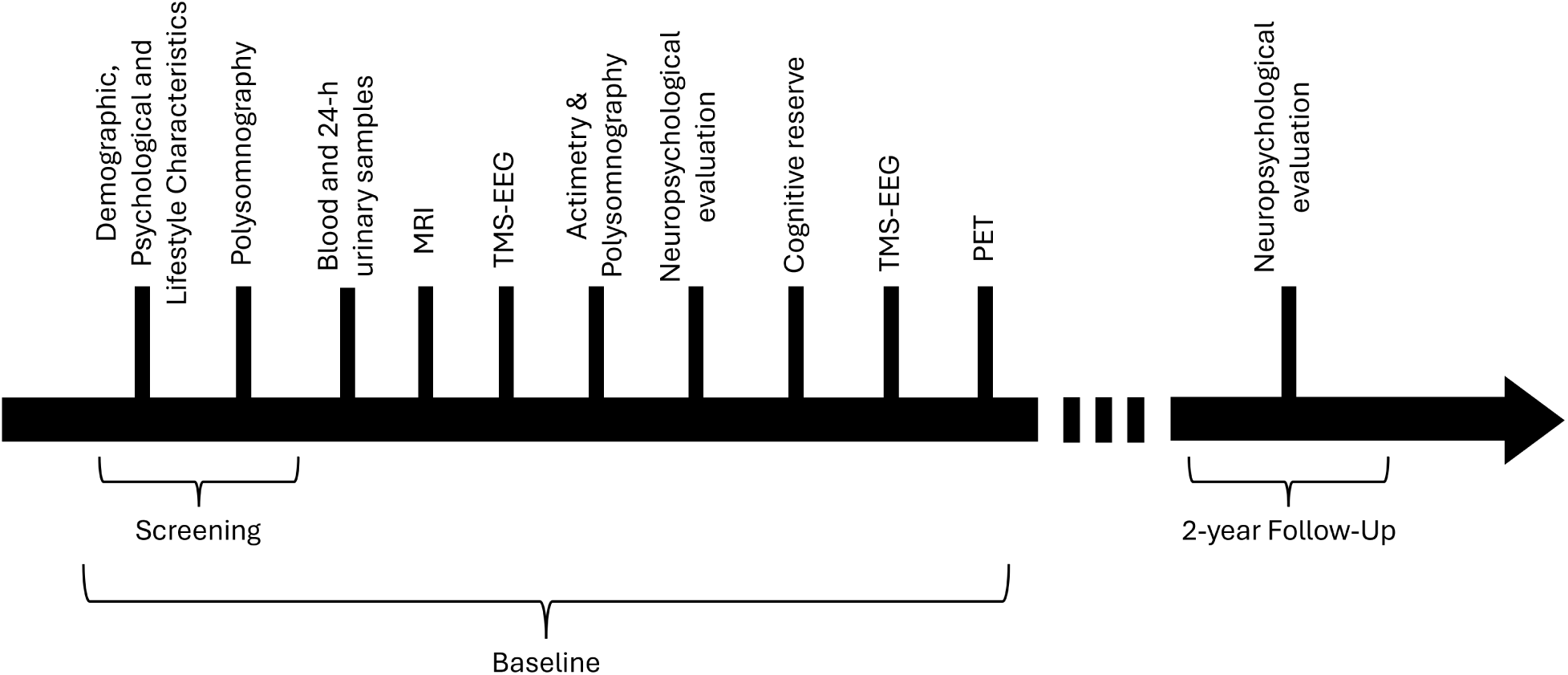
Timeline of data acquisitions. The baseline session was acquired between 2016 and 2018.

Baseline data collection consisted of multiple acquisition sessions aimed at capturing complementary aspects of brain and behavioral functioning. Blood samples were collected to perform genotyping of common single nucleotide polymorphisms, including *APOE* status. Neuroimaging data were acquired using a comprehensive MRI protocol combining high-resolution structural imaging, quantitative multi-parametric mapping, and multi-shell diffusion-weighted imaging. Molecular imaging was performed using positron emission tomography (PET). Amyloid-*β* burden was assessed in all participants using either [18F]Flutemetamol or [18F]Florbetapir radiotracers, while a subset of participants additionally underwent [18F]THK-5351 PET imaging to estimate tau-related processes. Cognitive and behavioral phenotyping included an extensive neuropsychological assessment administered over two sessions of approximately 75 minutes each, as well as self-report questionnaires assessing lifestyle factors contributing to cognitive reserve. This was followed by an in-laboratory overnight sleep assessment under polysomnography (PSG) to record sleep parameters.

Beyond these imaging and phenotypic data, participants completed electroencephalography (EEG) and combined EEG–transcranial magnetic stimulation (EEG–TMS) sessions to assess cortical excitability. Additional physiological samples (including saliva and urine) and derived measures (including cholesterol, glucose and melatonin levels), as well as actimetry data were also collected. These data are not included in the present release but will be subsequently made available. Similarly, the raw polysomnographic recordings (including EEG, ECG, EMG, and pupillometry), from which sleep parameters were derived, will be shared in a future data release. In addition to the two-year longitudinal cognitive assessment currently available, an ongoing seven-year follow-up evaluation has been completed in approximately 65 participants and will be incorporated into future releases of the dataset.

A longitudinal follow-up was conducted approximately two years after baseline acquisition in a subset of participants, focusing on reassessing cognitive performance using a targeted neuropsychological battery. This design enables the investigation of early cognitive trajectories and their relationship with baseline multimodal imaging and biomarkers.

### II.B. Participants

One hundred and one healthy late middle-aged (50-69 years old) French-speaking community-dwelling women and men (N = 101; 68 women [67.3%] and 33 men) enrolled in a study designed to identify biomarkers and lifestyle factors associated with normal cognitive ageing in the context of preclinical dementia. No participants reported any recent history of neurological or psychiatric disease or were taking medication affecting the central nervous system. All had normal or corrected-to-normal vision and hearing. Other exclusion criteria were sleep apnea/hypopnea index *≥* 15/h, assessed during an in-lab night of sleep under polysomnography (PSG), BMI *<* 18 and *>* 30 kg/*m*^2^, smoking, illicit drug consumption, sleep-medication use, excessive consumption of caffeine (*>* 7 cups/day) or alcohol (*>* 15 units/week), diabetes and shift-work over the past 6 months or transmeridian travel in the last 2 months.

Participants with high levels of depression and anxiety as assessed by the Beck Depression Inventory^26^ and by the Beck Anxiety Inventory^27^, respectively, were excluded (i.e., score *>* 20), as well as participants with ongoing pharmacological treatment of depression or anxiety. Participants with treated (*>* 6 months) hypertension and hypothyroidism were not excluded. All participants showed normal performance on the Mattis Dementia Rating Scale^28^ (i.e., score *>* 130).

The experimental procedures were approved by the local faculty-hospital ethic committee of the University of Liège and were in accordance with the Code of Ethics of the World Medical Association (Declaration of Helsinki) for experiments involving humans. All participants gave their signed informed consent before the experiment. They acknowledged that their data were fully anonymized and that their ethics consent included acceptance of anonymized data sharing. They also received financial compensation. Moreover, MRIs with contrast similar to those used in clinical practice were inspected by a neurologist who found no major anomalies, allowing to exclude other potential pathologies (vascular, tumor).

### II.C. Data Collection

All data from the COFITAGE dataset were acquired at the GIGA In Vivo Imaging platform^1^, University of Liège (Liège, Belgium), a multimodal neuroimaging facility dedicated to advanced in vivo brain imaging.

#### II.C.1. Magnetic Resonance Imaging (MRI)

All MRI data were acquired using a 3 Tesla MRI scanner (Siemens MAGNETOM Prisma, Siemens Healthineers, Erlangen, Germany) and were performed using a 64-channel head/neck coil. The standard MRI protocol incuded structural MRI, quantitative multiparametric MRI as well as multi-shell diffusion-weighted imaging.

Multi-parameter mapping (MPM) data were acquired using a 3D multi-echo fast low-angle shot (FLASH) sequence with RF spoiling^29^. The protocol comprised three co-localized multi-echo acquisitions with different contrast weightings: T1-weighted (T1w), proton density-weighted (PDw) and magnetization transfer-weighted (MTw). For the T1w acquisition, eight echoes were acquired with echo times ranging from 2.34 to 18.72 ms (effective TE = 14.04 ms), repetition time (TR) = 24.5 ms, and flip angle = 21*^◦^*. The PDw acquisition used the same echo train (TE range: 2.34-18.72 ms; effective TE = 4.68 ms) with TR = 24.5 ms and flip angle = 6*^◦^*. The MTw acquisition consisted of six echoes (TE range: 2.34-14.04 ms; effective TE = 2.34 ms), with TR = 24.5 ms, flip angle = 6*^◦^*, and an additional off-resonance magnetization transfer saturation pulse. The sequence was based on a 3D spoiled gradient-echo (FLASH) readout and identical geometric parameters were used across contrasts to allow subsequent quantitative map estimation.

T1-weighted (T1w) anatomical images were acquired using a 3D magnetization-prepared rapid gradient-echo (MPRAGE) sequence (TR = 1900 ms, TE = 2.19 ms, inversion time (TI) = 900 ms, flip angle = 9*^◦^*, acquisition matrix = 240 x 256 x 224, voxel size = 1 x 1 x 1 *mm*^3^). The acquisition was performed using a 3D inversion-recovery prepared spoiled gradient-echo readout, providing whole-brain coverage.

T2-weighted (T2w) images were acquired using a 2D turbo spin-echo (TSE) sequence (TR = 9240 ms, TE = 80 ms, flip angle = 180*^◦^*, acquisition matrix = 448 x 448 x 60, voxel size = 0.4 x 0.4 x 1.2 *mm*^3^). The acquisition was performed using a multi-slice 2D scheme with a spin-echo-based contrast. T2-weighted images were acquired in an oblique-coronal orientation perpendicular to the long axis of the hippocampus and positioned to cover its entire structure.

Diffusion-weighted images (DWI) were acquired using a single-shot spin-echo echoplanar imaging (EPI) sequence (TR = 7400 ms, TE = 69 ms, flip angle = 90*^◦^*, acquisition matrix = 96 × 106 x 70 axial slices, voxel size = 2 × 2 × 2 mm^3^). Fat suppression was applied, and diffusion sensitization was performed along multiple shells, including 15 directions at b = 650 s/mm^2^, 30 directions at b = 1, 000 s/mm^2^, and 60 directions at b = 2, 000 s/mm^2^. In addition, 13 non-diffusion-weighted (b = 0 s/mm^2^) volumes were acquired with reversed phase-encoding directions to enable susceptibility-induced distortion correction. Images were acquired with a phase-encoding direction of posterior-to-anterior (*j*-). The effective echo spacing was 0.31 ms, resulting in a total readout time of 32.4 ms. The acquisition was performed using a 2D multi-slice scheme.

Magnetic field maps (FMap) were acquired using a dual-echo gradient-echo sequence to estimate spatial variations in the static magnetic field (B_0_). The acquisition consisted of two echoes with echo times TE_1_ = 10 ms and TE_2_ = 12.46 ms and a flip angle of 90*^◦^*, using a 2D acquisition scheme. The phase difference between the two echoes was used to compute voxel-wise frequency offsets, providing a field map that characterizes B_0_ inhomogeneities across the brain. These field maps were subsequently used to correct susceptibility-induced geometric distortions in EPI data.

#### II.C.2. Positron Emission Tomography (PET)

Positron emission tomography (PET) data were acquired using an ECAT EXACT+ HR scanner (Siemens, Erlangen, Germany) providing molecular imaging of amyloid-*β* and tau-related processes. All participants (N = 101) received a single intravenous injection in an antecubital vein with a target activity of approximately 185 MBq. Among these 101 participants, 100 underwent amyloid PET imaging using either ^18^F-florbetapir (N = 3) or ^18^F-flutemetamol (N = 97), while 67 underwent tau PET imaging using ^18^F-THK5351. Overall, 66 participants had both amyloid and tau PET scans, 34 had only amyloid PET, and one participant had only tau PET.

For amyloid PET imaging, a late dynamic acquisition protocol was used. Scanning started approximately 85 minutes post-injection to ensure sufficient tracer uptake, and consisted of four 300 s frames (total duration *≈* 20 minutes), followed by a 10-minute transmission scan (Germanium-68) used for attenuation correction.

For tau PET imaging, a transmission scan (10 minutes) was first acquired, followed by a dynamic acquisition starting immediately after tracer injection. The dynamic protocol comprised 32 time frames spanning early (15 s) to late phases (up to 90 minutes post-injection), including both short (15–60 s) and longer frames (up to 300 s) to capture tracer kinetics. The total acquisition duration was approximately 100 minutes. Head motion was minimized using a thermoplastic mask to stabilize the participant’s head on the scanner bed. In addition, participants were instructed to remain still throughout the acquisition. All PET data were reconstructed using a filtered back-projection algorithm, with standard corrections applied for attenuation (transmission-based), scatter, random coincidences, and dead time.

#### II.C.3. Phenotype

In the COFITAGE dataset, a comprehensive and multidimensional assessment framework has been employed to characterize participants’ cognitive functioning, cognitive reserve, and psychological as well as subjective, circadian, and well-being profiles. A broad neuropsychological battery was administered to capture performance across major cognitive domains, including attention, executive functioning, memory, and global cognition. In parallel, demographic, psychological, and lifestyle characteristics, along with *APOE* genotype, and sleep parameters from polysomnography and actimetry, were collected. Cognitive tests were grouped into subcategories based on the primary cognitive function each test is designed to assess.

##### Cognition & Cognitive Reserve Assessment

Attention and processing speed were assessed using several complementary measures. The Choice Reaction Time task^30^ provided an index of basic attentional processes and psychomotor speed. The d2 Test of Attention^31^ evaluated selective and sustained attention under time pressure, while the Digit Symbol Substitution Test^32^ measured processing speed, visual scanning, and associative learning. In addition, Part A of the Trail Making Test (TMT-A)^33^ assessed visual attention and rapid sequencing abilities.

Executive functions were examined through tasks requiring inhibition, cognitive flexibility, and strategic retrieval. The Stroop Color-Word Test^34^ measured inhibitory control by requiring participants to suppress automatic reading responses. Part B of the Trail Making Test (TMT-B)^33^ assessed set-shifting and cognitive flexibility through alternating numeric and alphabetic sequencing. Verbal fluency tasks^35^, including phonemic and semantic fluency, evaluated generative search strategies and executive retrieval processes. The Raven’s Progressive Matrices^36^ provided an additional measure of executive control through abstract problem-solving and inductive reasoning.

Working memory was assessed using the forward and backward Digit Span task^32^, which measured short-term memory capacity and the manipulation of information, respectively. The visual n-back^37^ task also contributed to this domain by imposing dynamic working memory load.

Episodic memory was evaluated using multiple complementary measures. The Free and Cued Selective Reminding Test (FCSRT)^38^ assessed controlled encoding, storage and retrieval of verbal information. The Logical Memory task^39^ required immediate and delayed recall of short narratives, providing an index of narrative episodic memory. The Mnemonic Similarity Task (MST)^40^ measured pattern separation processes by requiring discrimination between highly similar stimuli, offering a sensitive index of hippocampal-dependent memory.

Language tasks were proposed as proxies of cognitive reserve: the Mill Hill Vocabulary Scale^41^, which measured receptive vocabulary and the French version of the National Adult Reading Test (NART)^42^, which estimated premorbid verbal intelligence through pronunciation of irregular words. Finally, global cognitive functioning was assessed using the Mini-Mental State Examination (MMSE)^41^ and the Mattis Dementia Rating Scale^28^, both of which provide broad indices of cognitive status across multiple domains.

Finally, cognitive reserve was also assessed using a computerized lifestyle questionnaire^43^ evaluating educational attainment, occupational complexity, physical activity and engagement in leisure activities across the lifespan.

##### Demographic, Psychological and Lifestyle Characteristics

Anxiety and depressive symptoms were assessed using the Beck Anxiety Inventory (BAI)^27^ and the Beck Depression Inventory (BDI)^26^, two 21-item self-report questionnaires measuring the severity of anxiety and depressive symptoms. Subjective cognitive complaints were evaluated with the 39-item Cognitive Difficulties Scale (CDS)^44^, in which participants rated the frequency of everyday cognitive problems over the past three weeks on a 0–4 Likert scale. The CDS is a sensitive measure of perceived cognitive difficulties, including in studies examining associations with A*β* burden. ^45^ ^46^ Daytime sleepiness was evaluated using the Epworth Daytime Sleepiness Scale^47^ and individual chronotype was determined using the Horne-Ö stberg questionnaire^48^, which assesses circadian preference for morningness or eveningness.

This dataset also includes core demographic, anthropometric and lifestyle information. Variables comprise age, sex, years of education, body weight, BMI and handedness. Lifestyle measures include self-reported weekly alcohol consumption and daily intake of coffee and tea.

##### *APOE* Genotyping

Genotyping was performed using DNA extracted from blood samples. Genome-wide single nucleotide polymorphisms (SNPs) were assessed using the Illumina Infinium OmniExpress-24 BeadChip (Illumina, San Diego, CA, USA), mapped to the human reference genome build hg19 (GRCh37). Genotype imputation was conducted using the Sanger Imputation Server^2^ with the Haplotype Reference Consortium (HRC, release 1.1)^49^ as the reference panel and Eagle2 for pre-phasing^50^.

##### Sleep Assessment

A first night of sleep was recorded at the laboratory under full polysomnography to avoid potential first night effects and exclude volunteers with parasomnia and sleep apnea (Apnea Hypopnea Index *≥*15 /hr). A second night of sleep was recorded with EEG, following 1 week of regular sleep-wake schedule based on each participant’s preferred bed and wake-up time^51^. The compliance was verified by actimetry (Actiwatch, Cambridge Neurotechnology, UK) and sleep diary. The timeline of the protocol, including the screening nights, is illustrated in Figure 1. Sleep was recorded with N7000 amplifiers (EMBLA, Natus, Planegg, Germany). The recording comprised 11 electroencephalography (EEG) derivations, placed according to the 10-20 system (F3, Fz, and F4; C3, Cz, and C4; P3, Pz, and P4; O1 and O2), two bipolar electrooculogram, and two bipolar submental electromyogram electrodes. Sampling was set at 200 Hz^52^ ^51^ ^53^.

### II.D. Data preprocessing

MR imaging data were converted from DICOM to NIfTI format using `spm dicom convert.m`, version 7136, SPM12.3 (Wellcome Centre for Human Neuroimaging, London, UK) implemented in MATLAB R2019b. Metadata were extracted during conversion and stored in accompanying JSON sidecar files. In addition, DWI-specific metadata required for BIDS compliance (including gradient information, `reconmatrixPE`, `effectiveEchoSpacing` and `TotalReaoutTime`) were extracted during the `dcm2niix` (v1.0.20250506) conversion and cross-checked using `dicom2nifti` (v2.6.2) conversion. The data were made BIDS-compliant after conversion through `qMRI-BIDS`^54^.

PET data were reconstructed using the scanner standard reconstruction software, based on the filtered back-projection algorithm, with standard corrections applied for attenation (transmission-based), dead time, random events, and scatter^55,56,57^. The reconstructed images consisted of 128 × 128 × 63 voxels, with a spatial resolution corresponding to approximately 2.57 × 2.57 × 2.43 mm^3^. Decay correction and calibration were applied using a dose calibration factor of approximately 9.52 × 10^6^. PET data were converted from ECAT format to NIfTI using the SPM conversion utility (SPM *→* PET *→* Convert *→* ECAT *→* NIfTI). This procedure generates a NIfTI image file accompanied by a minimal JSON sidecar containing basic metadata. Following conversion, the data were processed using `PET2BIDS`^58^ and `PET-BIDS`^59^.

Phenotypic and neuropsychological data were curated and harmonized using custom Python scripts (Python 3.11.5). Data cleaning included consistency checks, handling of missing values, and verification of variable ranges. Genetic variables were derived from genotyped polymorphisms: rs429358 and rs7412 were coded as allele pairs^60^, and the *APOE ɛ* genotype (*ɛ*2, *ɛ*3, *ɛ*4 alleles) was inferred from their combination. Participants were subsequently categorized as *ɛ*4 carriers (heterozygous or homozygous) or non-carriers, and an ordinal *APOE* -based risk score was computed to reflect increasing genetic risk associated with *ɛ*4 allele dosage^61^. In parallel, EEG signals were re-referenced to the mean of the two mastoids, and sleep stages were automatically scored in 30 s epochs using a validated algorithm (ASEEGA, Physip, Paris, France)^51,52,53^.

To enable consistent BIDS formatting across all data modalities, all datasets were organized and harmonized according to the Brain Imaging Data Structure (BIDS) specification^24^ using `Bidsme` (v1.11.0)^25^. Prior to release, all participants were fully anonymized. Structural MRI images were defaced using `spm deface.m` from SPM12 to remove facial features. In addition, all identifying information was removed from imaging headers and phenotypic files, and participants identifiers were harmonized across modalities to ensure consistency with BIDS naming conventions. To further protect privacy, acquisition dates were shifted within a *±*15-day window while preserving intra-subject temporal consistency.

### II.E. Data Validation

All data were collected using standardized protocols and quality-controlled procedures to ensure consistency across participants and modalities. Quality control procedures were applied at multiple stages of preprocessing. Imaging data were inspected for artifacts, motion and reconstruction errors. Subjects or volumes failing predefined quality criteria were flagged for review, and exclusion criteria were documented where applicable.

Neuropsychological testing was conducted under strictly standardized conditions to ensure consistency across participants. All testing sessions took place at a similar time of day, with identical lighting and the same computer equipment. Experimenters received dedicated training to ensure uniform administration of instructions and procedures, minimizing any variability related to the tester. Paper-and-pencil tests were digitized through scanning for secure storage and subsequent score verification. Participants’ verbal responses were audio-recorded to enable post hoc verification, with a subset independently reviewed by a second evaluator to ensure scoring reliability.

As a final validation step, the dataset was validated using the BIDS Validator (version 3.0.0-alpha.3)^62^, confirming compliance of all files, metadata structures, and directory organization with the current Brain Imaging Data Structure specifications.

For two participants (sub-070 and sub-071), the MT-weighted (MTw) acquisition was performed with a field-of-view orientation differing from the corresponding PDw and T1w acquisitions. Inspection of the original DICOM headers confirmed that this orientation discrepancy was already present in the source data and did not result from the DICOM-to-BIDS conversion process. Visual quality control indicated that image quality was otherwise preserved and that the acquisitions remained usable for subsequent processing. Users should nevertheless be aware of this acquisition-specific characteristic when performing analyses relying on precise inter-contrast alignment within the multi-parameter mapping protocol.

### II.F. Data format and usage notes

#### II.F.1. Data format

The COFITAGE dataset is openly hosted on a public EBRAINS repository https://. It follows the standard BIDS directory structure (see Figure 2), with one folder per participant (`sub-xxx`). When multiple time points are available, session-level subfolders (`ses-yyy`) are included, and session information for phenotypic data is recorded in the corresponding TSV files. Each session contains modality-specific subdirectories—anat/, dwi/, fmap/ and pet/. PET and MRI data are stored as NIfTI (.nii.gz) files with accompanying JSON metadata in their respective modality folders. Phenotypic data are provided as TSV files in the phenotype/ directory, with detailed variable descriptions in associated JSON files.

**Figure 2:**
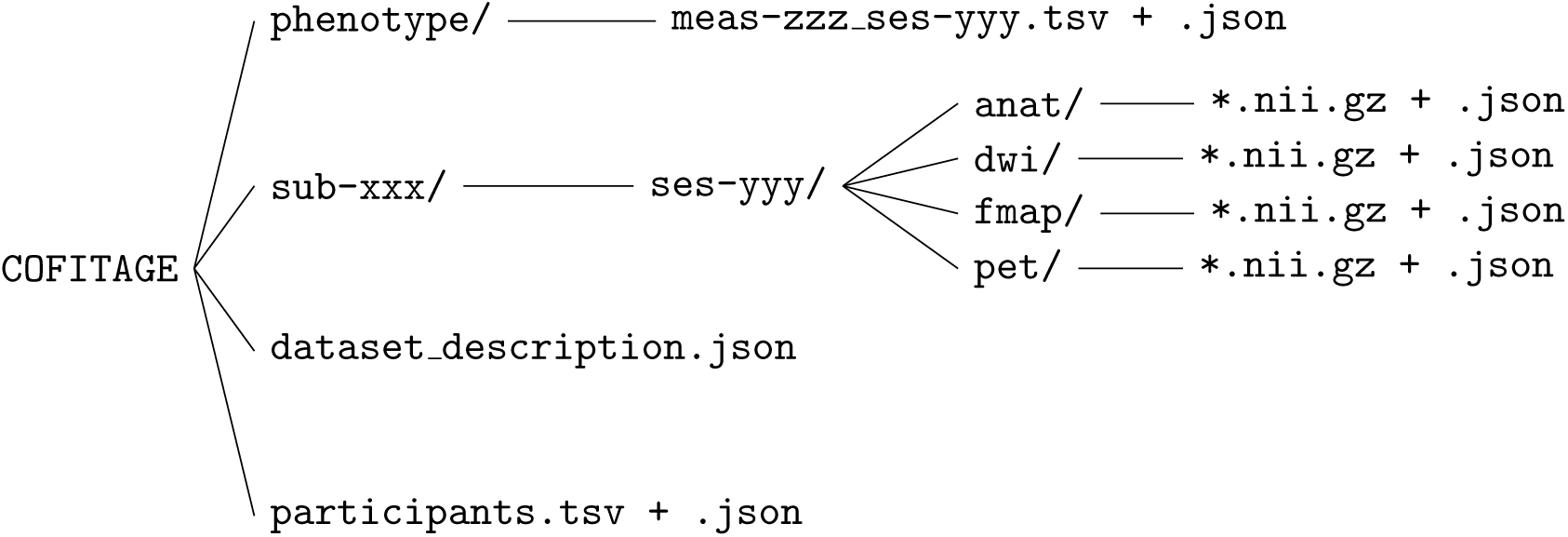
Overview of the dataset organization following the BIDS specification. The identifier `xxx` refers to the patient, `yyy` denotes the session and `zzz` specifies the measurement tool.

The top level of the dataset comprises three files: dataset description.json, which specifies general metadata about the dataset, including its name, authorship, and BIDS version; participants.tsv, which contains participant-level demographic and clinical variables; and participants.json, which provides the corresponding metadata describing the variables defined in participants.tsv.

#### II.F.2. Usage notes

Thanks to the BIDS organization of COFITAGE, users are encouraged to rely on the accompanying JSON sidecar files for detailed acquisition parameters and metadata associated with each imaging modality. Neuroimaging data include multiple contrasts and sequences; users should carefully select appropriate preprocessing pipelines depending on their research objectives (e.g., structural analysis, diffusion modeling, or quantitative MRI).

Phenotypic and neuropsychological data are stored as tab-separated values (TSV) files within the `phenotype/` directory, with detailed variable descriptions provided in accompanying JSON files. Users should pay particular attention to missing data patterns (n/a) and variable definitions when performing statistical analyses.

An overview of the datasets available per participant is provided in Table 2.

**Table 2:**
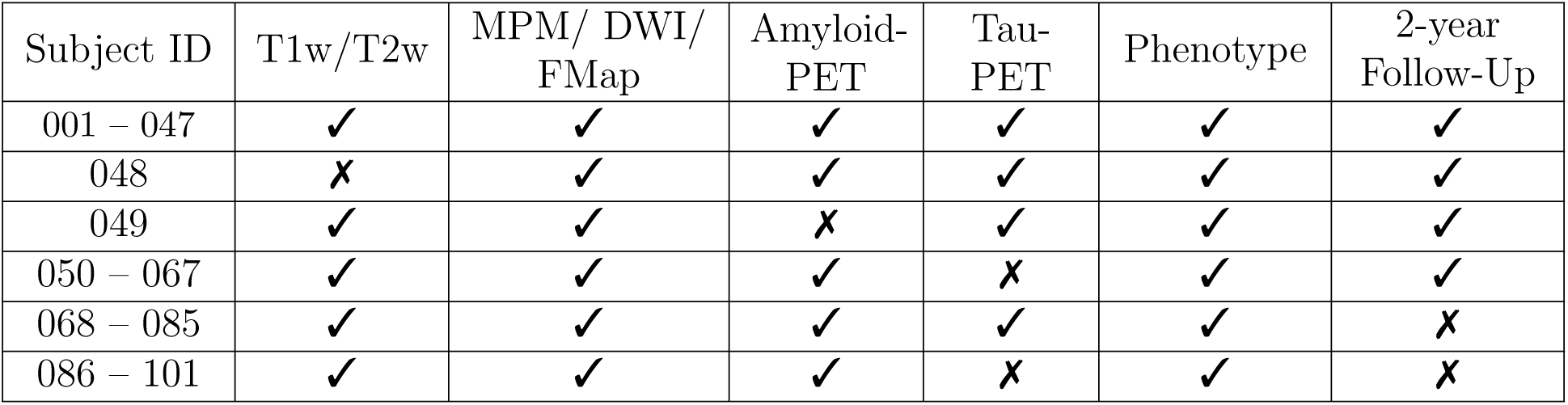
Overview of the data available per participant. The phenotype column comprises all phenotypic measures excluding genetic data.

## III. Discussion

Existing public neuroimaging datasets, such as Alzheimer’s Disease Neuroimaging Initiative (ADNI)^63^, Open Access Series of Imaging Studies (OASIS) ^64^, and Parkinson’s Progression Markers Initiative (PPMI)^65^, are valuable resources with large sample sizes and substantial statistical power. However, these initiatives are primarily disease-oriented, with protocols and outcome measures designed to support diagnosis and disease progression rather than the characterisation of normative cognitive aging. Their multicentric design and reliance on standardized clinical procedures often result in relatively coarse phenotyping, limiting the characterization of subtle inter-individual variability in brain and cognitive aging. For example, ADNI was initially established to validate biomarkers for Alzheimer’s disease (AD). Among its approximately 3400 participants, individuals aged 50–59 remain relatively under-represented, with only 210 subjects in this age range. Moreover, although ADNI includes several cognitive assessments, its phenotyping mainly targets diagnostic classification and disease staging rather than fine-grained quantitative and qualitative characterization of cognitive functioning.

In contrast, the COFITAGE cohort was specifically designed as a deeply phenotyped longitudinal multimodal study of healthy late middle-aged adults. Although smaller in sample size, it integrates neuroimaging, sleep physiology, genetics, and an extensive neuropsychological assessments within the same individuals, an approach rarely achieved in open-access datasets. Such rich phenotyping may enable the identification of biological and cognitive features associated with larger effect sizes and increased sensitivity to early aging-related alterations. Beyond discovery-driven analyses, the dataset provides a valuable framework to investigate mechanistic hypotheses generated, for example by linking the microstructural or molecular alterations to imaging findings derived from conventional MRI analysis.

Overall, studies based on the COFITAGE dataset converge toward a nuanced view of early cognitive aging. Rather than being primarily driven by overt AD pathology, interindividual differences in late-midlife cognitive performance appear to reflect an interplay between brain functional dynamics, sleep–wake regulation, and individual factors such as cognitive reserve and mental health^14,16,21^. The dataset supports a wide range of research applications, including studies of preclinical AD biomarkers, hippocampal vulnerability, amyloid-tau-metabolism interactions, sleep-dependent modulation of molecular pathology, microstructural correlates of cognitive reserve, and multimodal predictive modeling of cognitive trajectories. Its harmonized and open structure also makes it suitable for methodological developments in image processing, multimodal data fusion, machine learning, and reproducible neuroimaging pipelines.

In particular, cortical responsiveness and its modulation across wakefulness emerge as sensitive indicators of cognitive fitness^17,20,23^, while multiple findings highlight tight—yet complex and sometimes bidirectional—links between sleep characteristics and AD-related biomarkers^10,11,12^. Importantly, these relationships are not uniform, as some sleep features may be protective rather than detrimental^10^. In addition, dynamic processes such as day-today fluctuations in brain activity and perceived effort contribute to cognitive variability^19,22^, underscoring the importance of moving beyond static measures.

Moreover, several studies further emphasize the central role of individual differences, especially cognitive reserve and neuropsychiatric symptoms, which may outweigh biomarker effects at this stage^13,14,16^. Altogether, these findings suggest that early cognitive ageing arises from the complex interaction between functional, behavioral, and biological factors, positioning the COFITAGE dataset as a valuable resource for identifying early and potentially modifiable markers of cognitive decline.

Nevertheless, several limitations should be acknowledged. Although COFITAGE is a single-site cohort, which ensured greater consistency in data acquisition and neurophysiological assessments, this may limit the generalizability of the dataset to other acquisition settings. In addition, the sample size remains modest compared to large public cohorts, potentially limiting statistical power for subgroup and multimodal analyses. Finally, follow-up assessments are not available for all participants and did not include imaging data, which may restrict certain longitudinal and multimodal investigations.

Despite these limitations, the depth and multimodal richness of the dataset provide a unique opportunity to investigate early, subtle changes in ageing brain that are often inaccessible in large but less deeply phenotyped cohorts.

## IV. Conclusion

This work introduces a longitudinal, multimodal resource designed to characterize early brain ageing through the integration of neuroimaging, sleep, genetic, and cognitive assessments. It provides well-characterized foundation for investigating subtle brain changes associated with ageing and early neurodegenerative processes. In addition, this dataset addresses a critical gap in open-access resources by focussing on the 50-70 year age range, a key period during which early cognitive decline may emerge. By providing BIDS-standardized, quality-controlled, and openly accessible data, it enables multimodal analyses, supports interdisciplinary research, and facilitates the development of biomarkers and predictive models of brain aging.

## Footnotes

1 https://www.gigaivi.uliege.be/

